# First evidence of SARS-CoV-2 genome detection in zebra mussel (*Dreissena polymorpha*)

**DOI:** 10.1101/2021.05.28.446136

**Authors:** Antoine Le Guernic, Mélissa Palos Ladeiro, Nicolas Boudaud, Julie Do Nascimento, Christophe Gantzer, Jean-Christophe Inglard, Jean-Marie Mouchel, Cécile Pochet, Laurent Moulin, Vincent Rocher, Prunelle Waldman, Sébastien Wurtzer, Alain Geffard

## Abstract

The uses of bivalve molluscs in environmental biomonitoring have recently gained momentum due to their ability to indicate and concentrate human pathogenic microorganisms. In the context of the health crisis caused by the COVID-19 epidemic, the objective of this study was to determine if the SARS-CoV-2 ribonucleic acid genome can be detected in zebra mussels (*Dreissena polymorpha*) exposed to raw and treated urban wastewaters from two separate plants to support its interest as bioindicator of the SARS-CoV-2 genome contamination in water. The zebra mussels were exposed to treated wastewater through caging at the outlet of two plants located in France, as well as to raw wastewater at laboratory scale in controlled conditions. Within their digestive tissues, our results showed that SARS-CoV-2 genome was detected in zebra mussels, whether in raw and treated wastewaters. Moreover, the detection of the SARS-CoV-2 genome in such bivalve molluscans appeared even with low concentrations in raw wastewaters. This is the first detection of the SARS-CoV-2 genome in the tissues of a sentinel species exposed to raw and treated urban wastewaters. Despite the need for development for quantitative approaches, these results support the importance of such invertebrate organisms, especially zebra mussel, for the active surveillance of pathogenic microorganisms and their indicators in environmental waters.

**Graphical abstract:** 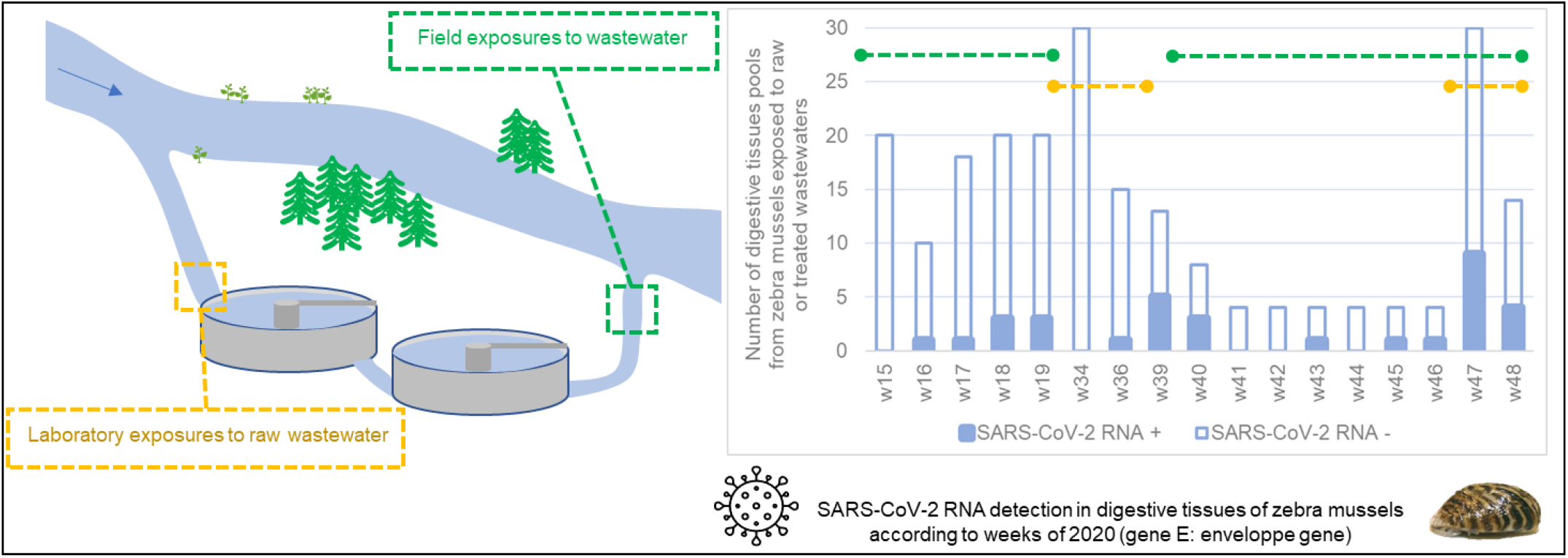

## 1. Introduction

Since several months, the world has been facing a historic viral pandemic. This pandemic was formalized by the World Health Organization (WHO) on March 11, 2020 and has a known origin in the Wuhan region in China (Langone et al., 2021; WHO, 2020a). Since then, this disease has spread around the world, causing many victims, and disrupting daily life. This pandemic (COVID-19) is caused by SARS-CoV-2, a coronavirus. The *Coronaviridae* includes seven virus species that infect humans, among them SARS-CoV and MERS-CoV, which appeared in the 2000s, and therefore SARS-CoV-2, discovered in December 2019 (Lu et al., 2020; Yang et al., 2020). In France, human contaminations were focused on two waves of contamination, from early March 2020 to mid-May 2020 and from mid-October 2020 to the end of 2020.

This virus is mainly transmitted by direct contact with an infected person or indirectly via infected droplets (Langone et al., 2021). These droplets are found in the air or on surfaces whose nature greatly varies the lifespan of the virus (Ren et al., 2020). However, since the SARS-COV-2 can infect and replicate both gastrointestinal glandular epithelial cells and respiratory system, the faecal-oral contamination cannot be excluded (Amirian, 2020; Heller et al., 2020; Peng et al., 2020). The occurrence of SARS-CoV-2 genome in the faeces is about 43% of COVID-19 patients and can longer be detected in digestive tract than in the respiratory one (Amirian, 2020; Kitajima et al., 2020; Zhang et al., 2021). This virus can therefore reach wastewater via sewages from cities and hospitals. The presence of SARS-CoV-2 genome has been detected in many raw wastewaters worlwide, especially during intense epidemiological phases (Balboa et al., 2020; Guerrero-Latorre et al., 2020; Kitajima et al., 2020; Nemudryi et al., 2020; Randazzo et al., 2020; Rimoldi et al., 2020; Wang et al., 2020; Wurtzer et al., 2020a). The treatments carried out in wastewater treatment plants (WWTPs) seem inactivate infectious SARS-CoV-2 since numerous studies attesting to its presence in the raw wastewater no longer observe it after biological treatments performed by some WWTPs (Balboa et al., 2020; Randazzo et al., 2020; Rimoldi et al., 2020; Singer and Wray, 2020). Nevertheless, some studies have detected SARS-CoV-2 genomes in treated wastewaters at wastewater treatment plants in France and Germany (Westhaus et al., 2021; Wurtzer et al., 2020b). This virus can also be detected in rivers in many developing countries, with rudimentary or in the absence of water treatment systems (Guerrero-Latorre et al., 2020), but also in developed countries (Polo et al., 2021; Rimoldi et al., 2020). However, there is still little knowledge concerning the survival of infectious SARS-CoV-2 in this aquatic environment. At a laboratory scale, Desdouits et al. (2021) demonstrated the accumulation of inactivated SARS-CoV-2 genome in different shellfish tissues of oysters (*Crassostrea gigas*). Wurtzer et al. (2021) have shown the presence of numerous forms of SARS-Cov-2 genomes in wastewaters including a small part of infectious and encapsidated particles using RT-qPCR and infectivity assays. Traces of SARS-CoV-2 genome were assessed in digestive tissues of *Ruditapes philippinarum* and *R. decussatus* taken from several coastal sites in Spain (Polo et al., 2021).

For several years, the use of sentinel species (i.e. bivalve molluscans) of the microbiological contamination of the environmental waters has intensified. The detection for many pathogens in filter-feeding and sessile organisms have many advantages and can complement the direct analyses of water matrices. Bivalve molluscans can indicate bacterial, protozoan or even viral contamination (Bighiu et al., 2019; Capizzi-Banas et al., 2021; Kerambrun et al., 2016; La Rosa et al., 2021). The high filtration capacity of bivalves allows them to filter large volumes of water (Palos Ladeiro et al., 2018; Polo et al., 2021). These invertebrate organisms can therefore be exposed to a panel of contaminants potentially more representative of their environment than that found in a water sample. Indeed, Bighiu et al. (2019) pointed to bacterial indicators 132 times higher in zebra mussels than in wastewater. Correlatively, the hepatitis A virus was detected in 16% of bivalve samples (*Mytilus galloprovincialis, Solen vagina, Venus gallina*, and *Donax trunculus*) against 9% in all water samples (La Rosa et al., 2021). This has also been demonstrated in zebra mussels for the Low Pathogenic Avian Influenza virus, but to a lesser extent (Stumpf et al., 2010). This filtration capacity is supplemented by an interesting bioaccumulation kinetics since the filter-feeding bivalves rapidly accumulate biological pollutants while being able to keep them several days (or even weeks) after the pressure in the environment has disappeared (Bighiu et al., 2019; Capizzi-Banas et al., 2021; Stumpf et al., 2010). This allows to have an integrative approach of water contamination over time. Also, the possibility to perform active biomonitoring through the caging allows a temporal and spatial assessment of the contamination, comparing different geographical sites or hydrosystems (Capizzi-Banas et al., 2021). Among indicator species, the zebra mussel, *Dreissena polymorpha*, has many advantages for biomonitoring programs under biological pressure (Kraak et al., 1991; Palos Ladeiro et al., 2014). This species is quite resistant to environmental pressures, is easy to handle and can be used in laboratory studies or in the field through the caging technique (Bervoets et al., 2005; Capizzi-Banas et al., 2021; Géba et al., 2020; Kerambrun et al., 2016; Le Guernic et al., 2020; Palos Ladeiro et al., 2018). The digestive tissues of bivalve molluscs is generally used to detect the presence of enteric viruses, especially enteroviruses, since it is the main site of contamination within the bivalve maybe due to specific receptors (Fuentes et al., 2014; Le Guyader et al., 2006; Lees and TAG, 2010; Suffredini et al., 2020). Desdouits et al. (2021) have reported accumulation of inactivated SARS-CoV-2 genome in digestive, mantle and gill tissues of oysters (*Crassostrea gigas*). This highlighted the potential interest of using digestive tissues of *D. polymorpha* for detecting SARS-CoV-2 genome in environmental waters.

In this context, the objectives of this study were: i) to know if SARS-CoV-2 genomes can be detected in zebra mussels at the inlet and / or at the outlet of two French wastewater treatment plants (WWTPs), namely Reims and center Seine (Ile-de-France public sanitation service, SIAAP), and ii), to determine if this organism can be used as a bioindicator of water contamination by this virus in field and laboratory exposures. These objectives are tested during the two waves of contamination observed in France.

## 2. Materials and methods

### 2.1. Zebra mussels

Zebra mussels (2.98 ± 0.38 g; 2.51 ± 0.19 cm) were collected during October and November of 2019 from Der lake (51290 Giffaumont-Champaubert, France, N 48°33’35”; E 4°45’11”) and brought back to the laboratory, where they were maintained in 1,000 L aerated tanks with 750 L of municipal drinking water (13.46 ± 1.77°C; pH 8.15 ± 0.17; 597 ± 27 mS/cm; 0.21 ± 0.05 mg/L nitrites; 58.05 ± 13.54 mg/L nitrates; 0.14 ± 0.42 mg/L ammoniac). Mussels were kept several months before the experiments, under these acclimation conditions. Throughout this acclimation step, mussels were fed *ad libitum*, twice per week, with *Nannochloropsis* (Nanno 3600, Planktovie, Marseille, France).

### 2.2. Reims and SIAAP WWTPs

The Reims WWTP is located at 16 chemin des Temples, 51370 Saint-Thierry (49°16’49.566” N, 3°59’32.625” E) and is managed by Grand Reims. The center Seine WWTP is located 5 Boulevard Louis Seguin, 92700 Colombes (48°55’57.936” N, 2°14’38.58” E) and is managed by the SIAAP. Their characteristics are summarized in Annex 1. Briefly, the two treatment plants have common characteristics, namely physical and chemical treatment of wastewater and sludge, as well as biological treatment of wastewater. The biological treatment of the two WWTPs is mechanical (anaerobic and aeration tanks, biofilters, etc.) and does not include a step with chlorine. At the end of the water treatment process, this water is discharged into the Vesle for the Reims WWTP, and into the Seine for that of the SIAAP.

### 2.3. Experimental designs

#### 2.3.1. *In situ* exposures to treated wastewaters

Two exposures to effluent were performed on dates corresponding to the two epidemiological waves observed in France. The first one was performed from 07^th^ April 2020 to 07^th^ May 2020, while the second one was performed from 25^th^ September 2020 to 27^th^ November 2020. These cages were deposited into the 1,000 L acclimation tanks. Polyethylene cages, having a volume of 931 cm^3^, and exhibiting 5 x 5 mm mesh, contained 150 mussels, and were then deposited by two at the study sites. For the site of Reims, cages were placed on the sediment with a water column height of at least 40 cm above them and were connected to the bank with a cable, while for the center Seine site, cages were placed inside the WWTP in a tank receiving treated wastewaters.

For the earlier experiment (April and May 2020), mussels were caged at the exit of the Reims WWTP (49°16’39.5” N, 3°59’06.5” E) and inside that of center Seine (48°55’57.936” N, 2°14’38.58” E). Caging and sampling kinetics are described in Table 1. The digestive glands of three zebra mussels were grouped together to have enough biological material for the analyses. At each sampling time, ten pools of three digestive glands are recovered for Reims WWTP, and five pools of three digestive glands for SIAAP WWTP. The samples were then directly frozen by liquid nitrogen vapours and then stored at −80°C before the analyses. Concerning the experiment conducted in October and November 2020, mussels were only exposed to the exit of Reims WWTP. As previously described, dissections were performed at laboratory and pools of digestive glands were then stocked at −80°C until RT-qPCR. For each different caging periods, less than 10% mortality was reported.

**Table 1:** Caging and sampling kinetics for exposures to treated wastewaters.

| Experiment | Locations | Start | Sampling times | End |
| --- | --- | --- | --- | --- |
| Spring 2020 | Reims WWTP | 07th April | D0 / D1 / D3 / D7 /<br>D14 / D21 / D28 | 05th May |
|  | Center Seine WWTP | 16th April | D4 / D7 / D11 / D14 /<br>D18 / D21 | 07th May |
| Autumn 2020 | Reims WWTP | 25th<br>September | D3 / D7 / D14 / D21 /<br>D28 / D35 / D41 /<br>D49 / D56 / D63 | 27th<br>November |

#### 2.3.2. Laboratory exposures to raw wastewater

Four laboratory experiments were performed on dates corresponding to the second epidemiological wave observed in France. The first one was performed from 18^th^ August to 22^nd^ August 2020, the second from 02^sd^ September to 05^th^ September 2020, the third exposure was realised from 24^th^ September to 27^th^ September 2020, while the fourth one was performed on one week from 16^th^ November 2020 to 23^rd^ November 2020.

The experimental procedure is identical for all four experiments, as described below. Before the experiments, mussels were placed in 10 L aerated glass tanks in the dark with control of the temperature at 13°C. Four tanks containing each 30 *D. polymorpha*, were implemented: i) with 100% (4 L) of Cristaline Aurele drinking water (spring Jandun, France); ii) 10% of raw wastewater coming from the WWTP of Reims and collected the day before (drinking water q.s. 4 L); ii) 33% of raw wastewater (drinking water q.s. 4 L); and iv) 100% of raw wastewater. These waters were changed every day, and the input of raw wastewater came, each experiment day, from a sample the day before. Concerning the first exposure (August 2020), samples were collected on D1, D2, D3 and D4. For both September exposures, samples were collected only on D3, and mussels were not fed during the experimentation step, while concerning the last exposure (November 2020), that lasted longer (sampling time on D1 and D7), mussels were fed every day with Nanno 3600 algae (Planktovie, Marseille, France) before the water change. For this last exposure, two tanks containing 30 zebra mussels were placed for the 100% raw wastewater condition. As previously described, dissections were performed at laboratory and pools of digestive glands were then stocked at −80°C before RT-qPCR analysis. During the exposures carried out at the end of September and in November, mussels in 100% and 33% raw sewage conditions could be dissected respectively before D3 and D7 according to their general condition (in particular the time required to close the valves). For these experiments, 15 pools of 3 mussels were dissected before D3 (September) or D7 (November) because of the toxicity of untreated wastewater, undiluted or two-thirds diluted.

### 2.4. SARS-CoV-2 genome detection in wastewater

Analyses of SARS-CoV-2 genome in raw wastewater were realised by the Obepine group (Réseau Obepine, 2021). Briefly, virus particles were concentrated by ultracentrifugation of 11 mL of wastewater sample and RNA genome were extracted according to Wurtzer et al. (2020a). SARS-CoV-2 genes RdRp (RNA-dependent RNA polymerase), E (envelope protein) and N (nucleocapsid protein) were assessed and quantified by RT-qPCR according to Pasteur Institute protocol (WHO, 2020b), Corman et al. (2020) and CDC protocol (U.S. Department of Health and Human Services, 2020) respectively (Table 2). Then these data were synthesized into an indicator obtained by data assimilation with a digital model of the Kalman filter type (Forward-Backward). This graph was constructed only with envelop protein gene. Data for the Reims and center Seine (SIAAP) WWTPs were collected from April 2020 to January 2021, and compared to periods of confinement and curfew observed in France (Réseau Obepine, 2021). This information is available on the Obepine network site (Réseau Obepine, 2021).

**Table 2:**
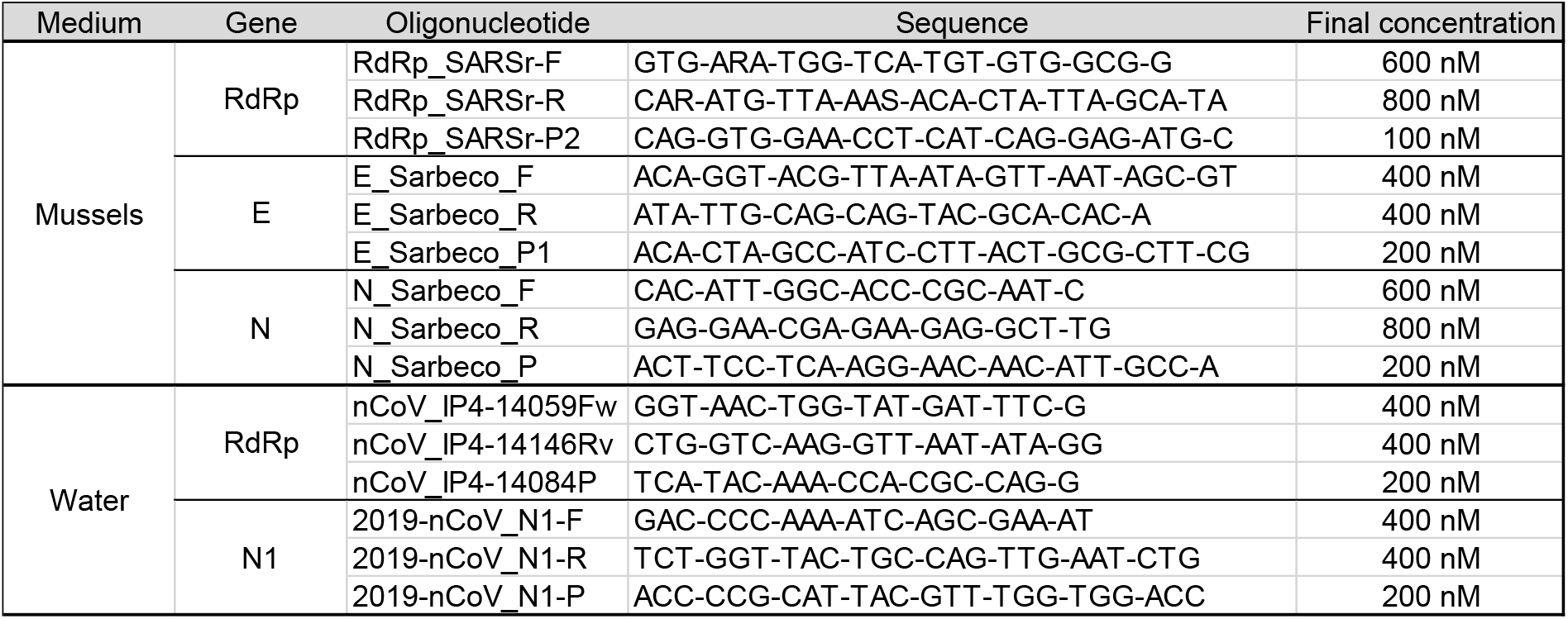
List and characteristics of primers (F and R) and probes (P) used for Rt-qPCR analyses. From Corman et al. (2020). E: envelope protein gene; RdRp: RNA-dependent RNA polymerase gene; N or N1 (in wastewater): nucleocapsid protein gene.

### 2.5. SARS-COV-2 GENOME detection in digestive tissues of D. polymorpha

#### 2.5.1. RNA quantification

After thawing of samples, 200 μL of proteinase K for 200 mg of digestive tissues was added (3 U/ml, Euromedex, Souffelweyersheim, France). Samples were then homogenized several seconds with an ultra-turrax (Ika-Werk, Janke & Kunkel, Staufen im Breisgau, Germany). Then, cells were lysed by adding trizol reagent and the whole was vortexed (Molecular Research Center Inc., OH, USA). Chloroform (VWR) was added and vortexed 30 seconds with samples and then incubated 15 minutes at room temperature. The aqueous phase containing the nucleic material was recovered after centrifugation (12,000 g, 15 min, 4°C). The following steps of RNA extraction were realised using the PureLink™ RNA mini kit (Invitrogen, ThermoFisher Scientific, MA, USA) following the manufacturer recommendations, until recovering RNA in RNAse free water. RNA samples were frozen (−20°C) until reverse transcription polymerase chain reaction.

#### 2.5.2. Reverse transcription polymerase chain reaction and RNA detection

SARS-CoV-2 RNA detection was based on works of Corman et al. (2020), and performed with SuperScript™ III one-step RT-PCR with platinum™ Taq (Invitrogen). Genes tested in this article were: RdRP: RNA-dependent RNA polymerase gene; E, an envelope protein gene and N, nucleocapsid protein gene. Primers and probes used come from the study of Corman et al. (2020), were provided by Eurogentec (Liege, Belgium) and are described below (Table 2 Table 2). Unlike water samples, the viral load within the digestive gland mash cannot be preconcentrated. Characteristics of RT-qPCR were: 10 min at 55°C (RT) / 3 min at 95°C / 50 cycles of 15 sec at 95°C / 30 sec at 58°C (CFX96 Touch Real-Time PCR System, BioRad, CA, USA). NTC controls were realised by adding molecular-grade water, positive controls were performed by adding SARS-CoV-2 positive control (COV019 batch number 20033001, Exact Diagnostics, TX, USA) before RT-qPCR, and extraction controls were performed by adding 10 μL of this positive standard to digestive gland pools from mussels not exposed (between dissection and freezing). This positive extraction control allowed the obtention of an extraction yield between initial and final concentration of 70% for the E and N genes, and of 28% for the RdRp gene. The positive detections of the SARS-CoV-2 genome in the digestive tissues of zebra mussels were validated by a second passage of these samples in reverse transcription polymerase chain reaction.

## 3. Results and discussion

### 3.1. Detection of SARS-CoV-2 genomes in raw wastewaters

Obepine group has perfomed the wastewater analyses on the two WWTPs studied, and summarized Figure 1A (Réseau Obepine, 2021). Table 3 contains the concentrations of the three targeted SARS-CoV-2 genes in raw wastewaters. These data were averaged over the week for caging exposure to treated wastewater, or over the duration of exposure during laboratory exposures to raw wastewater. The contamination profiles of untreated wastewater by SARS-CoV-2 from Reims and the center Seine WWTPs in 2020 were remarkably similar, and wastewater from both sites exhibited concentrations of comparable values (Table 3). During the first exposures of zebra mussels to treated wastewater (spring), water contamination by the SARS-CoV-2 genome was very high (almost 500,000 copies/L for E gene), but dropped considerably until it reached its lowest values at the end of these exposures (Figure 1A or <DL, Table 3). On the other hand, the exposures to treated wastewater carried out at the end of 2020 corresponded to a period when the index was quite high (between 50 and 150, *Figure 1*). During this second caging exposure, genome concentrations of SARS-CoV-2 in raw water remained stable (between 38,000 and 91,000 copies/L for E gene, Table 3). This range of values was also found within exposures carried out in the laboratory after half of September 2020. Indeed, a notable increase in concentrations between the two experiments carried out in September 2020 was observed (Table 3).

**Figure 1:**
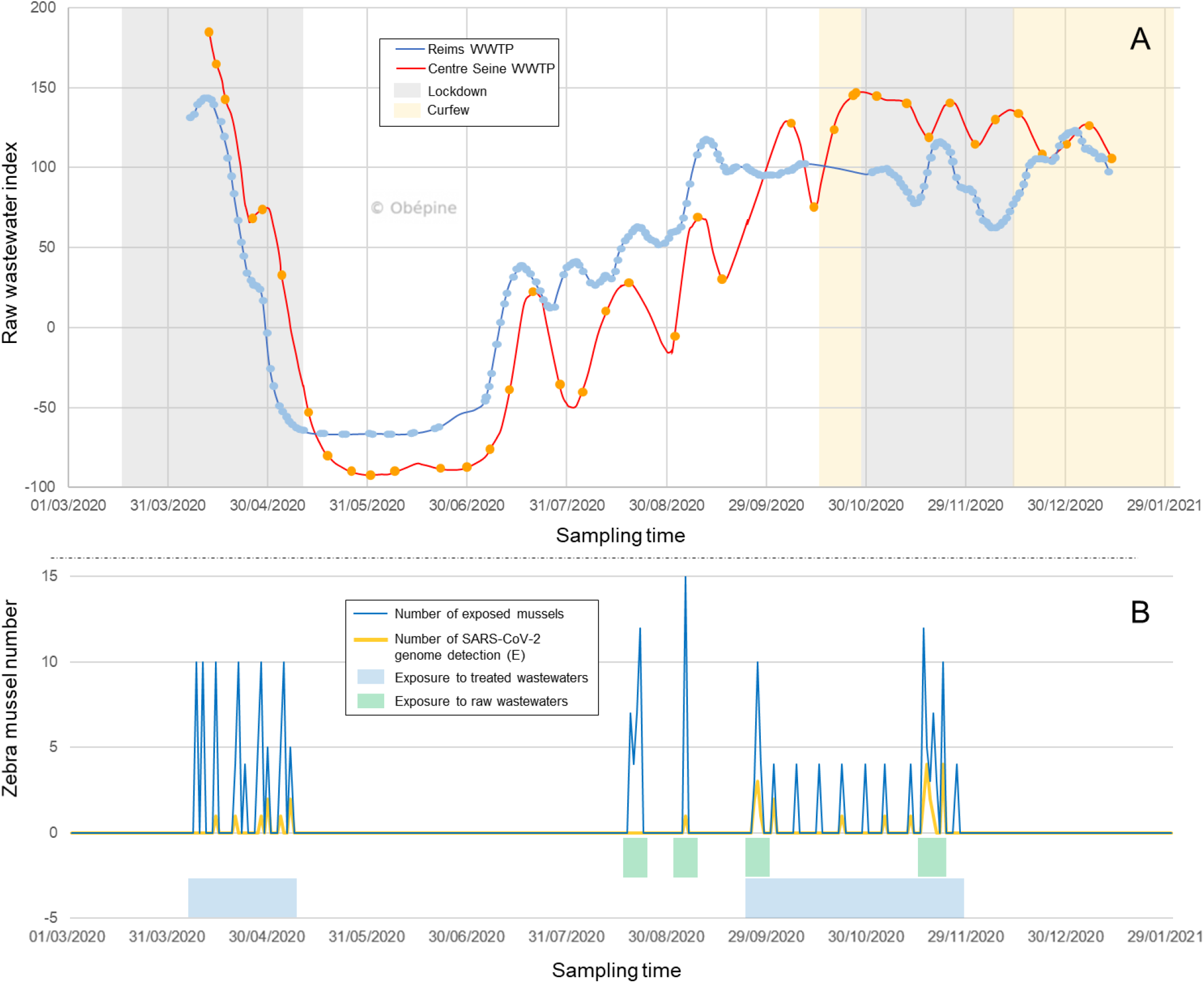
Detection of E gene of SARS-CoV-2 in raw wastewater (A) and in pool of digestive glands of zebra mussels (B) from the March 1^st^ 2020 to January, 29^th^ 2021. A: raw wastewater index of SARS-CoV-2 genome (gene E) from Reims (blue) and center Seine WWTPs (orange), according to OBEPINE group. Data are represented as a trend index based on RT-qPCR quantification on the E gene of the SARS-CoV-2 genome and assessed with a digital model of Kalman filter type (Forward-Backward). Confinement and curfew periods for Reims city were indicated by different colors. B: number of pools of digestive glands of zebra mussels exposed to raw wastewater or caged at the exit of Reims and center Seine WWTPs (blue curve) and number of pools with detection of at least one SARS-CoV-2 gene (orange curve). The periods of exposure to affluents or effluents from WWTPs are represented by rectangles of different colors.

**Table 3:** Concentrations (gene copies/L) of the SARS-CoV-2 genome in raw wastewater from WWTPs in Reims and center Seine, averaged over the week or over the duration of exposure. The concentrations under the various dilution conditions are estimates. Data are expressed as mean ± standard deviation (SD). The concentration estimate for the dilution conditions were obtained with respect to the 100% condition. E: envelope protein gene;RdRp: RNA-dependent RNA polymerase gene; N1: nucleocapsid protein gene; NA: not analysed; DL: detection limit.

| Experiment | Condition / Week | RNA concentration in raw wastewater<br>(average over the week or over the duration of the experiment) |  |  |  |  |  |  |  |  |
| --- | --- | --- | --- | --- | --- | --- | --- | --- | --- | --- |
|  |  | *RdRp* gene (copies/L) |  |  | *E* gene (copies/L) |  |  | *N1* gene (copies/L) |  |  |
|  |  | Mean | ± | SD | Mean | ± | SD | Mean | ± | SD |
| Spring caging<br>(April-May 2020) | Reims WWTP (W15) | NA | ± | NA | 489525 | ± | 411855 | NA | ± | NA |
|  | Reims WWTP (W16) | NA | ± | NA | 464844 | ± | 426420 | NA | ± | NA |
|  | Reims WWTP (W17) | NA | ± | NA | 21891 | ± | 17883 | NA | ± | NA |
|  | Reims WWTP (W18) | 398 | ± | 406 | < DL | ± | < DL | NA | ± | NA |
|  | Reims WWTP (W19) | 390 | ± | 202 | 20634 | ± |  | NA | ± | NA |
|  | center Seine WWTP (W16) | NA | ± | NA | 223704 | ± | 16364 | NA | ± | NA |
|  | center Seine WWTP (W17) | NA | ± | NA | NA | ± | NA | NA | ± | NA |
|  | center Seine WWTP (W18) | NA | ± | NA | 44178 | ± |  | NA | ± | NA |
|  | center Seine WWTP (W19) | NA | ± | NA | 2045 | ± |  | NA | ± | NA |
| Autumn caging<br>(September-October-November 2020) | Reims WWTP (W39) | 39276 | ± | 23004 | 80507 | ± | 39936 | 141385 | ± | 189176 |
|  | Reims WWTP (W40) | 14983 | ± | 12229 | 71795 | ± | 50457 | 31469 | ± | 19527 |
|  | Reims WWTP (W41) | 12110 | ± | 6586 | 77802 | ± | 55418 | 32343 | ± | 15197 |
|  | Reims WWTP (W42) | 5110 | ± | 2640 | 37909 | ± | 22827 | 19212 | ± | 15800 |
|  | Reims WWTP (W43) | 8169 | ± | 5701 | 41674 | ± | 19787 | 21552 | ± | 7286 |
|  | Reims WWTP (W44) | 11287 | ± | 2161 | 59840 | ± | 19573 | 36114 | ± | 21821 |
|  | Reims WWTP (W45) | 12417 | ± | 8961 | 73853 | ± | 29035 | 42215 | ± | 16489 |
|  | Reims WWTP (W46) | 8388 | ± | 2969 | 40722 | ± | 18186 | 46769 | ± | 39254 |
|  | Reims WWTP (W47) | 9156 | ± | 3901 | 79050 | ± | 78280 | 27520 | ± | 19756 |
|  | Reims WWTP (W48) | 3814 | ± | 4826 | 91106 | ± | 124838 | 15137 | ± | 11540 |
| 1st laboratory<br>exposure (August<br>2020) | 100 % raw wastewater | 6173 | ± | 2942 | 21400 | ± | 12297 | 24157 | ± | 25522 |
|  | 33 % raw wastewater | 2037 | ± | 971 | 7062 | ± | 4058 | 7972 | ± | 8422 |
|  | 10 % raw wastewater | 617 | ± | 294 | 2140 | ± | 1230 | 2416 | ± | 2552 |
| 2nd laboratory<br>exposure<br>(September 2020) | 100 % raw wastewater | 4244 | ± | 421 | 17237 | ± | 8801 | 30611 | ± | 18872 |
|  | 33 % raw wastewater | 1401 | ± | 139 | 5688 | ± | 2904 | 10102 | ± | 6228 |
|  | 10 % raw wastewater | 424 | ± | 42 | 1724 | ± | 880 | 3061 | ± | 1887 |
| 3rd laboratory<br>exposure<br>(September 2020) | 100 % raw wastewater | 32230 | ± | 19355 | 73026 | ± | 41874 | 57763 | ± | 33155 |
|  | 33 % raw wastewater | 10636 | ± | 6387 | 24098 | ± | 13818 | 19062 | ± | 10941 |
|  | 10 % raw wastewater | 3223 | ± | 1935 | 7303 | ± | 4187 | 5776 | ± | 3316 |
| 4th laboratory<br>exposure<br>(November 2020) | 100 % raw wastewater | 9913 | ± | 4086 | 113242 | ± | 120853 | 26232 | ± | 18650 |
|  | 33 % raw wastewater | 3271 | ± | 1348 | 37370 | ± | 39882 | 8657 | ± | 6154 |
|  | 10 % raw wastewater | 991 | ± | 409 | 11324 | ± | 12085 | 2623 | ± | 1865 |

### 3.2. Detection of SARS-CoV-2 genome in digestive tissues of zebra mussels exposed to treated wastewaters

The number of pools of digestive tissues from mussels caged in potentially contaminated wastewater as well as the number of pools with detection of the SARS-CoV-2 genome (at least one of the three genes tested) are shown on Figure 1B and on Figure 2A. Table 4 described the detections of the SARS-CoV-2 genome in zebra mussel samples.

**Figure 2:**
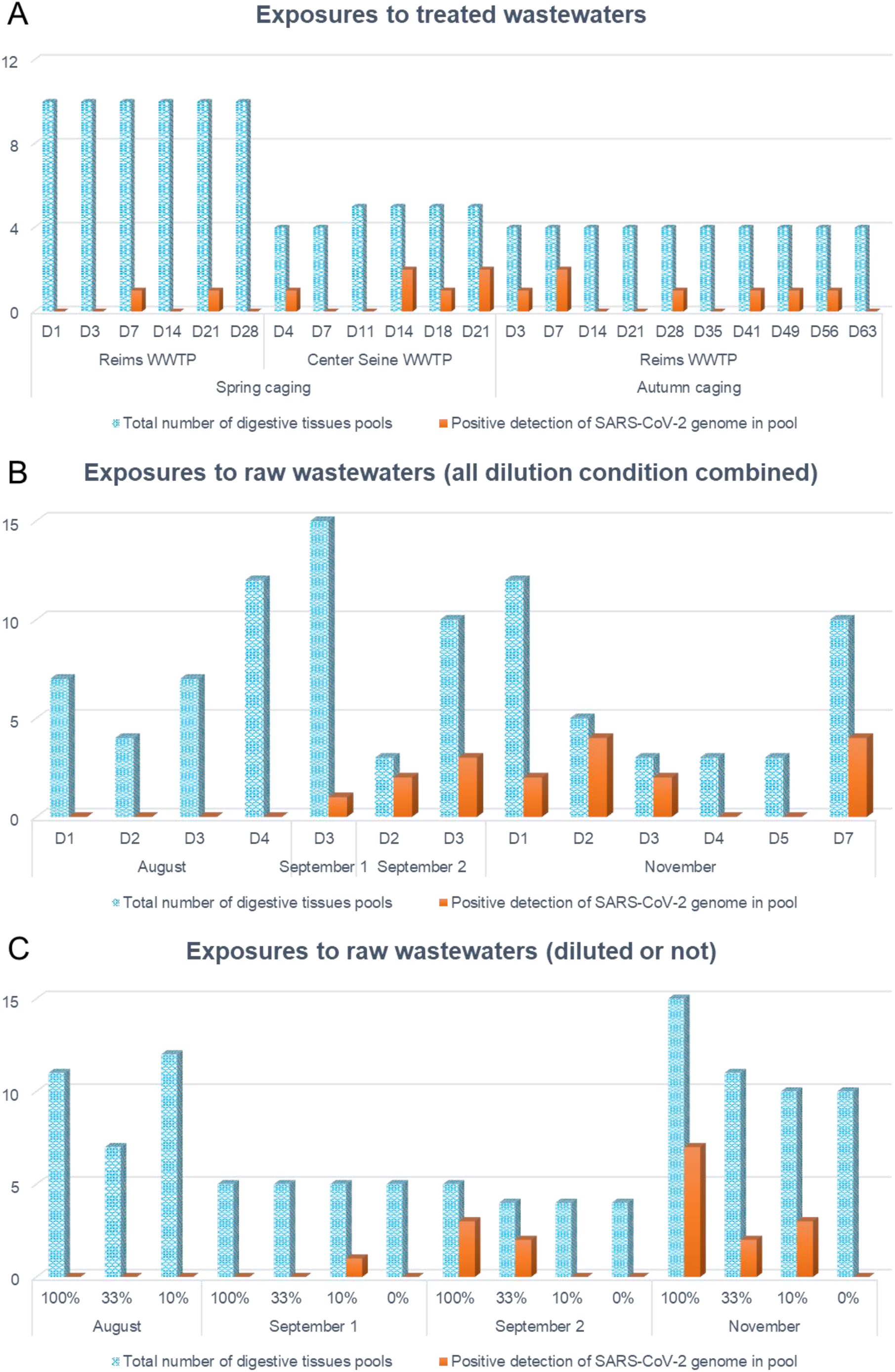
Total number of digestive tissues pool exposed (blue) and number of positive detection of SARS-CoV-2 genome in pools (orange) according to exposure, exposure condition and sampling times. A: Results obtained after exposure to treated wastewaters (spring and autumn) on the zebra mussels caged after Reims and center Seine WWTPs according to sampling times. B: Results obtained after exposure to raw wastewaters (August, September 1 and 2 and November exposures) from Reims WWTP according to sampling times (all dilution conditions combined). C: Results obtained after exposure to raw wastewaters (August, September 1 and 2 and November exposures) from Reims WWTP according to experiment and dilution conditions.

**Table 4:** Presence (+) or absence (-) of SARS-CoV-2 RNA in digestive glands of zebra mussels according to tested genes and exposure conditions. The samples presented in this table are positive for at least one of the three genes tested. E: envelope protein gene; RdRp: RNA-dependent RNA polymerase gene; N: nucleocapsid protein gene.

| Experiment | Date | Exposure time | Exposure condition | RdRp gene detection | E gene detection | N gene detection |
| --- | --- | --- | --- | --- | --- | --- |
| Spring caging | April 14, 2020 | D7 | Reims WWTP | - | + | - |
| Spring caging | April 20, 2020 | D4 | SIAAP WWTP | - | + | - |
| Spring caging | April 28, 2020 | D21 | Reims WWTP | - | + | - |
| Spring caging | April 30, 2019 | D14 | SIAAP WWTP | - | + | - |
| Spring caging | April 30, 2020 | D14 | SIAAP WWTP | - | + | - |
| Spring caging | May 04, 2020 | D18 | SIAAP WWTP | - | + | - |
| Spring caging | May 07, 2020 | D21 | SIAAP WWTP | - | + | - |
| Spring caging | May 07, 2020 | D21 | SIAAP WWTP | - | + | - |
| 2nd laboratory exp. | Sept. 05, 2020 | D3 | 10% raw wastewater | - | + | - |
| 3rd laboratory exp. | Sept. 26, 2020 | D2 | 100% raw wastewater | - | + | + |
| 3rd laboratory exp. | Sept. 26, 2020 | D2 | 100% raw wastewater | - | + | - |
| 3rd laboratory exp. | Sept. 27, 2020 | D3 | 100% raw wastewater | - | + | - |
| 3rd laboratory exp. | Sept. 27, 2020 | D3 | 33% raw wastewater | - | + | + |
| 3rd laboratory exp. | Sept. 27, 2020 | D3 | 33% raw wastewater | - | + | + |
| Autumn caging | Sept. 28, 2020 | D3 | Reims WWTP | - | + | + |
| Autumn caging | Oct. 02, 2020 | D7 | Reims WWTP | - | + | - |
| Autumn caging | Oct. 02, 2020 | D7 | Reims WWTP | - | + | - |
| Autumn caging | Oct. 23, 2020 | D28 | Reims WWTP | - | + | - |
| Autumn caging | Nov. 05, 2019 | D41 | Reims WWTP | - | + | - |
| Autumn caging | Nov. 13, 2020 | D49 | Reims WWTP | - | + | - |
| Autumn caging | Nov. 20, 2020 | D56 | Reims WWTP | - | + | - |
| 4th laboratory exp. | Nov. 17, 2020 | D1 | 100% raw wastewater | - | + | + |
| 4th laboratory exp. | Nov. 17, 2020 | D1 | 33% raw wastewater | - | + | - |
| 4th laboratory exp. | Nov. 18, 2020 | D2 | 100% raw wastewater | - | + | - |
| 4th laboratory exp. | Nov. 18, 2020 | D2 | 100% raw wastewater | - | + | - |
| 4th laboratory exp. | Nov. 18, 2020 | D2 | 100% raw wastewater | - | + | - |
| 4th laboratory exp. | Nov. 18, 2020 | D2 | 100% raw wastewater | - | + | - |
| 4th laboratory exp. | Nov. 19, 2020 | D3 | 100% raw wastewater | - | + | + |
| 4th laboratory exp. | Nov. 19, 2020 | D3 | 100% raw wastewater | - | + | - |
| 4th laboratory exp. | Nov. 23, 2020 | D7 | 33% raw wastewater | - | + | - |
| 4th laboratory exp. | Nov. 23, 2020 | D7 | 10% raw wastewater | - | + | - |
| 4th laboratory exp. | Nov. 23, 2020 | D7 | 10% raw wastewater | - | + | - |
| 4th laboratory exp. | Nov. 23, 2020 | D7 | 10% raw wastewater | - | + | - |

The first objective of our study was whether the SARS-CoV-2 genome could be detected by the zebra mussel caged at the exit of the WWTP. RNA of SARS-CoV-2 was found in digestive glands of mussels caged at the exit of both center Seine and Reims WWTPs (Table 4, Figure 2A). These detections covered, for each season, the entire exposure period (from April 14 to May 07, 2020 during spring caging, and from September 18 to November 20, 2020 during autumn caging, Table 4). Since the concentration values of the various SARS-CoV-2 genes were obtained in raw wastewater, the connection with their detection in caged mussels exposed to treated wastewater must be considered with caution. During the first caging campaign, corresponding to a decreasing phase of the raw wastewater index (Figure 1A), 10% of the exposed mussel pools showed positivity to the SARS-CoV-2 genomes in their digestive tissues (Figure 2A). Surprisingly, when only the results from the center Seine WWTP were considered, this percentage raised to 21%, compared to 5% at the outlet of Reims WWTP. In Reims, the two positive samples were reported in week 16, corresponding to very high concentrations of viral genomes in raw wastewater (464,844 copies of E gene per liter), but also in week 18, during which however the concentrations in the wastewater were below the detection limit (Table 3). For the center Seine WWTP, even with less data, the same observation was made, namely that the detection of SARS-CoV-2 genome in mussels mainly occurred during weeks 18 and 19 when the concentrations found in the raw wastewater were much lower (44,178 and 2,045 copies of E gene per liter respectively, Table 3). The detection of the SARS-CoV-2 genomes in digestive tissues of zebra mussels was therefore possible even with small amount present in the aquatic environment, and this detection lasted several days.

The second experiment was performed when the concentration of the SARS-CoV-2 genome in raw wastewater increased until a plateau (Figure 1A), and showed 18% of positivity to the virus genome in mussels (only for the Reims WWTP, Figure 2A). Looking more closely, genome positivity for SARS-CoV-2 in mussels was mainly observed during weeks when the concentration in the water was quite high (about 70,000 copies of E gene per liter, weeks 40, 45, 47, especially for the E gene, Table 3).

Several other studies have assessed the presence of SARS-CoV-2 genomes upstream and downstream of WWTPs. Most of them observed presence of the viral genomes in raw wastewater from urban WWTP (Balboa et al., 2020; Randazzo et al., 2020; Rimoldi et al., 2020). All these studies have reported the absence of SARS-CoV-2 genome in treated wastewaters after secondary ± tertiary treatments. Wurtzer et al. (2020b) and Westhaus et al. (2021) have nonetheless detected SARS-CoV-2 genomes after WWTP in France and Germany, respectively. Correlatively, Guerrero-Latorre et al. (2020) and Rimoldi et al. (2020) have reported presence of SARS-CoV-2 genome in rivers not linked to water treatment plant. There are therefore still gray areas as to the fate of this virus within hydrosystems.

Among the three genes used to detect SARS-CoV-2 genome, only the envelope (E) and of the nucleocapsid (N) genes were detected in mussels (Table 4). Even with maximum concentration of the samples, no detection of the RNA-dependent RNA polymerase (RdRp) gene was reported, contrary to N gene detected only with the maximum concentration (Table 4). The same conclusion can be made with regard to the concentrations of the three SARS-CoV-2 genes in untreated wastewater. In fact, the concentrations for the E gene were approximately 6 times higher than those of the RdRp gene and 2 times higher than those of the N gene (Table 3). These differences may be due to the various PCR efficiencies for these genes but also to the non-homogeneous fragmentation of viral genomes inside our biological matrix (Wurtzer et al., 2021). These discrepancies had already been revealed by other studies, for analyses of the SARS-CoV-2 genome in wastewater or in sludges. Several genes can be targeted by RT-qPCR to study the presence of the SARS-CoV-2 genome, based on the genes of envelope, nucleocapsid, ORF1ab, or even the RNA-dependent RNA polymerase (Kitajima et al., 2020). However, to date, there is no harmonization of procedures or standardization of the detection of SARS-CoV-2 genome and variations of results according to these assays were reported (Farkas et al., 2020). Corman et al. (2020) reported that the RdRp gene had a lower detection limit than the N and E genes. However, other studies observed different results, synthesized by Nalla et al. (2020). These authors have tested seven RT-qPCR assays linked to SARS-CoV-2 and concluded that N2 set and E gene are the most sensitive (Kitajima et al., 2020; Nalla et al., 2020), while Shirato et al. (2020) reported that only the RT-qPCR assays carried on the nucleocapsid gene worked for them. Rimoldi et al. (2020) evaluated the presence of 3 genes of the SARS-CoV-2 (Orf1ab, N, E) in different aquatic environments (WWTPs and rivers). Only one of the sites showed positivity to the SARS-CoV-2 genome with all 3 genes detected, and this is the only site where the E and N genes were both detected. Desdouits et al. (2021) used Corman’s E set for the envelope gene and IP4 set for RdRp gene, and these two genes were expressed in tissues lysates of *Crassostrea gigas* after controlled exposure to heat-inactivated SARS-CoV-2. In our study, the viral genome positivity of the digestive gland samples was mainly linked to the E gene, and a few of these samples also had positivity via the N gene (Table 4). Despite the lack of harmonization on the methods used, it would have been interesting to use other specific genes of SARS-CoV-2 (Orf1ab, RdRp IP4 set, other regions of N or E genes, etc.) to potentially improve its detection within digestive glands of zebra mussels.

Few studies had reported detection of SARS-CoV-2 genome in treated wastewaters. In our study, detection of the SARS-CoV-2 genome in the digestive glands of zebra mussels exposed at the WWTP outlet was observed. The use of a filter feeder and sessile species could explain this difference. The detection of SARS-CoV-2 genomes directly in wastewater was often represented by a value at a time point as well as on a volume of water which is not fully representative of the water mass. Correlatively, zebra mussels, because of their sessility and their filtration capacity, allow a more extensive characterization of the pollution of their environment (Kraak et al., 1991; Palos Ladeiro et al., 2014). In fact, these organisms can bioaccumulate biological and chemical pollutants for several days or even weeks, allowing pollution to be monitored over time, and filter a significant volume of water that is better representative of the mass of water (Bervoets et al., 2005; Palos Ladeiro et al., 2018; Wiesner et al., 2001). Concerning the SARS-CoV-2 genome, oysters have already proven their effectiveness by accumulating this virus during laboratory exposures (Desdouits et al., 2021). Also in the marine environment, the *Ruditapes* genus had shown its efficiency in accumulating the SARS-CoV-2 genome in their digestive tissues (Polo et al., 2021). The authors of this study concluded that mollusc bivalves can be used as biomonitoring tools for various anthropogenic contaminants, including the SARS-CoV-2 virus. During our study, zebra mussels were useful to detect SARS-CoV-2 genome in both untreated and treated wastewaters, even if the concentrations in wastewater was under the detection limit (1,000 copies/L). All these characteristics make such bivalve, and particularly zebra mussels, good indicators for the detection of SARS-CoV-2 genomes in such environments. These organisms can potentially support or even improve the sensitivity of the direct detection of the viral genome in water samples.

To further support the use of this sentinel species as an indicator of the presence of the SARS-CoV-2 genome in environmental waters, improvements on the extraction and detection of the SARS-CoV-2 genome in the digestive tissues of zebra mussels are required. Indeed, to improve detection of SARS-CoV-2 genome inside this complex biological matrix, the PCR cycle number has been increased to 50. Of the total positive detection data on E gene (Table 4), 73% had Cq lower than 42.75, but a few were higher (all Cq were comprised between 35.64 to 46.32). These high values underlined the limits of detection or extraction of this viral genetic material, and particularly in the biological matrix used here. Contrary to the quantification of SARS-CoV-2 genome in water, a pre-concentration step to concentrate the viral genome before analyses is not necessary (Kitajima et al., 2020). Moreover, the number of digestive glands per pool was not elevated (3). Several modifications can be considered to improve the viral extraction, such an addition of a purification step to limit as much as possible the enzymatic inhibitors which could be found in the biological matrix, preventing the good progress of the detection. In parallel, new experiments could be performed to improve the sensitivity of the detection of SARS-CoV-2 genome in mussels but also to characterize the bioaccumulation pattern in the tissues of *D. polymorpha*. These experiments must be performed in the laboratory in controlled conditions, to observe (or not) a dose-dependent accumulation relationship, and using untreated wastewater with higher levels of SARS-CoV-2 genomes than in treated water.

### 3.3. Zebra mussels as biological indicators of water contamination by the SARS-CoV-2 genome

The second aim of this study was to assess the interest of using zebra mussels as bioindicator of water contamination by the SARS-CoV-2 genome. Controlled laboratory exposures were therefore put in place to address the questions raised during exposure to treated wastewater. Also, to maintain the natural contamination by SARS-CoV-2 in the receiving aquatic environment, mussels were exposed to raw wastewater from Reims WWTP.

Regarding the experiments performed at a laboratory scale, the genome detection of SARS-CoV-2 genome was higher when the mussels were directly submitted to raw wastewater (100%) compared to the diluted wastewaters (33 or 10%, Figure 2C). Indeed, the most concentrated condition (100% raw wastewater) resulted in a positivity of 28% of the samples (10/32), greater than 15% (4/27) and 13% (4/31), caused respectively by conditions 33% and 10% of raw wastewater ratios. Zebra mussels were therefore useful to detect the SARS-CoV-2 genome in accordance with its presence in wastewaters. Furthermore, there was a similarity between the growing phase of the COVID-19 pandemic after summer 2020, confirmed by an increase of the raw wastewater index between July and October (*Figure 1*A), and the detection of the SARS-CoV-2 genome in mussels (Figure 2B). Indeed, when exposed to raw wastewaters in August, none of the 15 pools of exposed mussel digestive gland had the genome of this virus, and this number increased with time. During the first exposure in early September 2020, 7% (1/15) of the pools exhibited positivity for the SARS-CoV-2 genome, to increase to 38% (5/13) at the end of September, almost identical to the values found in November (33%, 12/36, *Figure 2*B). This result was in accordance with the sudden increase in the concentration of the SARS-CoV-2 genome in raw wastewater between early and late September (Table 3). This increase was all the more important for the E gene and also continued after September. For illustration, the concentrations of these genes in the raw wastewater were lower than those of the 33% diluted wastewater (Table 3). This originally suggested that the zebra mussel can be used as indicator of the SARS-CoV-2 genome detection in proportion to the contamination load present in freshwater environment and contributes to emphasis the uses of zebra mussels as sentinel species for SARS-CoV-2 contamination of wastewaters. Desdouits et al. (2021) and Polo et al. (2021) demonstrated the accumulation of anthropogenic virus by bivalves in several coastal sites including SARS-CoV-2 virus within digestive tissue. Contrary to Polo et al. (2021), Desdouits et al. (2021) but did not report the presence of SARS-CoV-2 genome in the field (in water or in bivalve molluscans). Nonetheless, even if the laboratory experiments allowed to expose zebra mussels to higher SARS-CoV-2 genome contamination, the experimental plan used in our study (short exposure due to the possible toxicity of raw wastewaters) only allowed the qualitative detection of the presence of SARS-CoV-2 genome in organisms but did not allow the genome quantification. Indeed, the possible toxicity of raw wastewater, causing an advanced dissection of organisms exposed to the most concentrated raw sewage conditions (18% of samples during the exposure at the end of September and 26% of samples during the last exposure, in November), did not allow mussels to be exposed any longer. To dispense with the toxicity of raw wastewater, a longer exposure of the mussels (from 14 to 21 days) in the laboratory to a non-infectious SARS-CoV-2 or to low pathogenic CoV strains could improve characterization of virus accumulation within mussels (Desdouits et al., 2021; Wurtzer et al., 2020b).

Thanks to our results, the use of this bivalve as a bioindicator and possible matrix to follow the presence of SARS-CoV-2 genome in water is conceivable, whether at the level of treatment plants, but also at the level of freshwater (rivers, etc.) or in countries whose water treatment structures are still underdeveloped. As announced by several recent studies, the bivalve taxon represents a complementarity, even a more than plausible alternative for the detection of viruses in the environment (Capizzi-Banas et al., 2021; Desdouits et al., 2021; La Rosa et al., 2021; Polo et al., 2021). Various fields of application can therefore be envisaged, such as environmental biomonitoring for health purposes.

## 4. Conclusion

Out of a total of 666 mussels exposed to water potentially contaminated by the SARS-CoV-2 genome, i.e. 222 pools of digestive glands, 33 pools showed positivity to the genome of this virus, representing almost 7%. This detection was observed during the two major epidemiological phases in France and both with raw wastewater and treated wastewater. Moreover, SARS-CoV-2 genomes was detected in *D. polymorpha* as well at the outlet of the Reims wastewater treatment plant as that of center Seine one. This corroborated the results in untreated wastewater but also brought a novelty with the resilience of the genetic material of the virus after treatment of these waters. This detection is proportional to the contamination in the wastewaters and can allow a temporal and spatial monitoring. The zebra mussel therefore appears to be an attractive candidate for detecting the presence of the SARS-CoV-2 genome in raw and treated wastewaters, but also in other hydrosystems. The detection of the genome of other enteric viruses could be relevant using this sentinel species.

## Supporting information

Annex 1

## Acknowledgments

The authors are deeply grateful to the MeSeine observatory from SIAAP (Ile-de-France public sanitation service) and the Obepine group for their contribution and advice to this study. This study was also supported by the ACTIA VIROcontrol Joint Technological Unit.

## Notes

### Competing Interest Statement

The authors have declared no competing interest.

